# Freestanding hydrogel lumens for modeling blood vessels and vasodilation

**DOI:** 10.1101/2021.10.19.464875

**Authors:** Ashley M. Dostie, Hannah G. Lea, Ulri N. Lee, Tammi L. van Neel, Erwin Berthier, Ashleigh B. Theberge

**Author notes:** Denotes co-first author (Ashley M. Dostie, Hannah G. Lea, Ulri N. Lee).

## Abstract

Lumen structures exist throughout the human body, and the vessels of the circulatory system are essential for carrying nutrients and oxygen and regulating inflammation. Vasodilation, the widening of the blood vessel lumen, is important to the immune response as it increases blood flow to a site of inflammation, raises local temperature, and enables optimal immune system function. A common method for studying vasodilation uses excised vessels from animals; major drawbacks include heterogeneity in vessel shape and size, time-consuming procedures, sacrificing animals, and differences between animal and human biology. We have developed a simple, user-friendly *in vitro* method to form freestanding cell-laden hydrogel rings from collagen and quantitatively measure the effects of vasodilators on ring size. The hydrogel rings are composed of collagen I and can be laden with human vascular smooth muscle cells, a major cellular and structural component of blood vessels, or lined with endothelial cells in the lumen. The methods presented include a 3D printed device (which is amenable to future fabrication by injection molding) and commercially available components (e.g., Teflon tubing or a syringe) to form hydrogel rings between 2.6-4.6 mm outer diameter and 0.79-1.0 mm inner diameter. Here we demonstrate a significant difference in ring area in the presence of a known vasodilator, fasudil (*p*<0.0001). Our method is easy to implement and provides a foundation for a medium-throughput solution to generating vessel model structures for future investigations of the fundamental mechanisms of vasodilation (e.g., studying uncharacterized endogenous molecules that may have vasoactivity) and testing vasoactive drugs.

## Introduction

Lumen structures exist throughout the body including in the glandular organs such as the breast and prostate, and blood vessels of the circulatory system. Vasodilation, the widening of the blood vessel lumen, is important to the immune response as it increases blood flow to a site of inflammation, raises local temperature, and enables optimal immune system function.^1^ Over the last two decades, tissue engineers and cell biologists have been working towards performing cell culture experiments in a three-dimensional environment, as opposed to two-dimensional culture.^2–4^ It is well accepted that in the human body, cells are encapsulated in a 3D environment (extracellular matrix, ECM) and receive signals very differently than they would in a 2D polystyrene culture plate. Incorporation of relevant hydrogels, multi-culture of different cell types, or generation of specific architectures, such as tubular structures, have resulted in changes in cell behavior that more closely recapitulate *in vivo* cell function and morphology.^5, 6^ Such observations have been increasingly helpful in drug discovery and tissue engineering due to increased cell to cell contact, cell communication, and cell-ECM interactions.^3^ Current methods for studying vasodilation and constriction involve time-consuming *ex vivo* methods utilizing blood vessels excised from animals (e.g., rabbit and rodent models), and simplified *in vitro* methods that capture a portion of the biological response.^7–10^ While excised animal vessels are valuable in understanding cell signaling in an *ex vivo* environment, procedures are time intensive, requiring processing of the excised tissue and then almost immediate testing of samples; additionally, the responses observed in these excised vessels may not be representative of a response in human tissue.^11^ An innovative model system is needed that is (1) simple to use and multiplex, enabling rapid adoption by biology laboratories, (2) incorporates multiple human cell types, and (3) can quantify the degree of dilation.

To offer an alternative to animal methods and clinical trials in humans,^12–14^ several groups have recently developed *in vitro* assays with primary human smooth muscle cells to study the effects of vasoactive compounds.^11, 15–18^ Alford *et al*. created what they called ‘muscular thin filament’, or MTFs, which are strips of polydimethylsiloxane (PDMS) with adhered vascular smooth muscle cells.^15^ The MTFs were imaged and analyzed, measuring constriction or dilation by curvature of the MTF.^15, 17, 19^ Alternatively, several groups have performed vasoactivity tests on tissue engineered blood vessels (TEBVs) as part of characterization of TEBVs to be used for drug testing^16^ or to be potentially implanted as vascular grafts.^18^ Recently, Tseng *et al*. developed a platform to study vasoactivity of vascular smooth muscle cells that were 3D printed into a ring configuration that fit within the well of a 96-well plate.^11^ Other well-based methods have been created that enable cells to settle and self-assemble into a ring formation.^20, 21^ We sought to add to this body of work by developing a user-friendly method for generating blood vessel mimics that require only commercially available supplies and simple parts produced with an inexpensive resin 3D printer. Additionally, the device can be scaled up by injection molding in polystyrene, a common cell culture material.^22^

Here, we present a modular casting method to form a 3D hydrogel structure with a lumen (i.e., a ring) that is embedded with smooth muscle cells. The method is user friendly and amenable to medium-throughput experiments (~100 rings per day). These hydrogel rings are 2.6-4.6 mm in outer diameter, 0.79-1.0 mm in inner diameter, and are free-standing and transferable between well plates. The lack of attachment to any surface enables the hydrogel to be remodeled by the cells; thus, the addition of vasoactive compounds results in a visible and quantifiable readout. Our method enables the use of smooth muscle cells embedded in the hydrogel and endothelial cells in the lining of the lumen. Our method has the potential to be developed further and used to assay potential drug candidates to reduce harmful vasodilation in patients with various inflammatory diseases (e.g., asthma, rheumatoid arthritis, Raynaud’s syndrome).

## Results

Our goal was to develop a user-friendly protocol for producing cell-laden hydrogel rings that can be used for medium-throughput biological and drug screening experiments. Lumen structures are commonplace in the human body; our area of focus in this work was modeling blood vessels to study vasodilation. Here we present two casting methods to produce cell-laden hydrogel rings; our methods are easy to implement with 3D printed components and low-cost commercially available materials (e.g., Teflon tubing). A table comparing the general features of excised animal tissue models, standard closed microfluidic lumen models, and our freestanding method is included in the supplemental information (Table S1).

To use cell-laden hydrogel rings in vasoactivity experiments, it is important that the hydrogel is not attached to stiff materials that may prevent hydrogel deformation (i.e., dilation or constriction); therefore, our approach was to cast hydrogels using an easily removable mold. We used a simple 3D printed design and Teflon tubing to mold precursor hydrogel solution into a ring that could be removed and placed in a standard 96-well plate. Current 3D printed molds consist of 6 posts per device, enabling the generation of 6 hydrogel rings per mold. It is feasible to make ~100 rings per day with each rack taking ~10 minutes total for assembly and loading of hydrogel. Incubation time varies on the hydrogel used (1 h for collagen). Other methods that utilize specialized assemblies to hold or rotate the rings during incubation limit the throughput of hydrogel ring structures to approximately <10 per experiment. Additionally, testing vasodilation with *ex vivo* vessels from mice or rabbits and wire myography only enables the measurement of 1-4 vessels at a time.^23^ A PDMS spacer was applied to the device, and Teflon tubing was added onto each post (Figure 1Aii). The PDMS spacer holds up the Teflon tubing and enables the tubing to slide down the post and unmold the hydrogel ring when the spacer is removed. F127 (a polyethylene glycol (PEG)-based hydrogel) was added to seal any gaps between the tubing and the 3D printed post. We chose F127 as a sacrificial layer because it is not toxic to cells and easily dissolves once placed in a liquid such as cell culture media or phosphate buffered saline (PBS). After the F127 gelled, collagen I was added to form the ring. Once the collagen I polymerized the PDMS spacer was removed and the device was placed in PBS. The tubing was pushed down into the gap created by the removal of the PDMS spacer to reveal the hydrogel rings which were then released into solution; the PBS dissolved the sacrificial F127 (Figure 1Aii). Collagen laden with primary human umbilical artery smooth muscle cells (HUASMCs) was used to form cell-laden hydrogel rings. A live/dead stain was performed with calcein AM and ethidium homodimer-1, showing the cells maintained excellent viability (Figure 1Bi). Further, arrays of hydrogel rings can efficiently be made with this method (Figure 1Bii).

**Figure 1.**
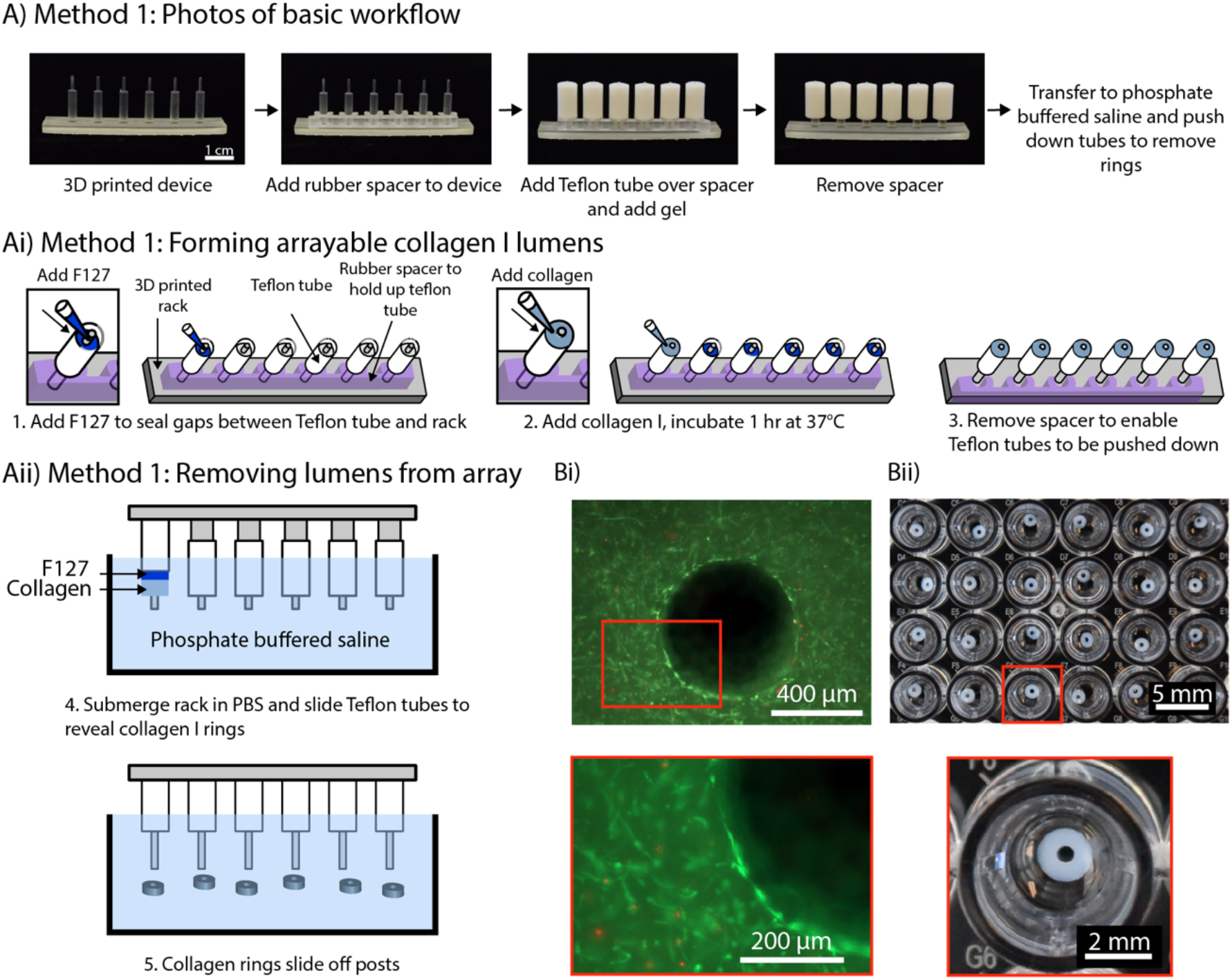
An arrayable method for fabricating cell-embedded free-standing collagen I lumens. (Ai) Photographs of device setup and basic workflow for Method 1. (Aii) A rubber spacer was added to the 3D printed device to hold up the Teflon tubes on each post. F127 hydrogel seals small gaps so the collagen I does not leak through. (Aiii) The rack was submerged in phosphate buffered saline, and the tubes were pushed down to remove the collagen rings and dissolve F127. (Bi) Fluorescence image of primary human umbilical artery smooth muscle cells embedded in a collagen I ring. Cells are stained with calcein AM (live, green) and ethidium-homodimer I (dead, red). (Bii) Image of an array of 3 mm OD and 1 mm ID hydrogel rings in a 96-well plate.

We evaluated the reproducibility of this method in two separate experiments with two devices of the same design (12 total collagen rings per experiment) in each experiment. While the rings can fit in a 96-well plate as shown in Figure 1B, the light from the imaging set up used to measure the rings reflects off the bottom of the concave meniscus, causing a glare over the collagen ring and impeding proper imaging. In a larger well plate, such as the 24 well plate, a glare was still present when imaging; however due to the larger diameter of the well relative to the collagen ring, the glare does not cover the ring (Figure 2A). This is important because when measuring the ring with ImageJ, the glare and white color of the collagen ring appear the same color which prevents the imaging software from differentiating the two areas.

**Figure 2.**
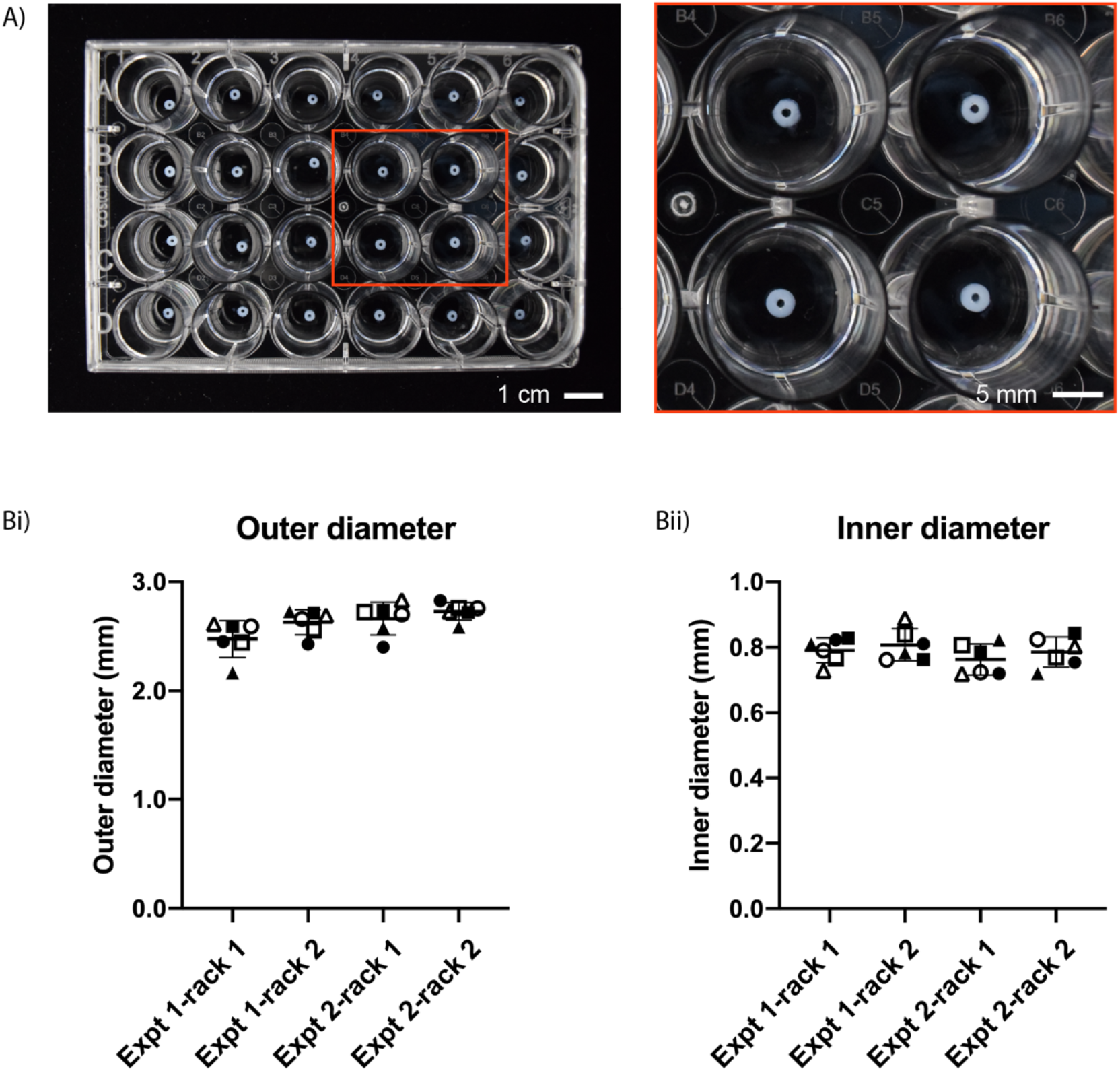
Reproducibility of collagen I rings designed with an outer diameter of 3.0 mm and inner diameter of 1.0 mm. (A) Collagen rings of equal size in a 24-well plate. (Bi) The average outer diameter from each device was 2.48±0.17 mm, 2.63±0.12 mm, 2.66±0.15 mm, 2.73±0.08 mm and (Bii) their respective inner diameters were 0.79±0.04 mm, 0.81±0.05 mm, 0.76±0.05 mm, and 0.79±0.05 mm. Each symbol pairs the OD measurements to ID measurements across Bi and Bii. Results are plotted from two independent experiments, each with two racks (one rack is an array of 6 lumens). Error bars are mean ± SD. Full set of data available in Table S2.

The designed dimensions for the collagen rings were 3 mm outer diameter and 1 mm inner diameter. Overall, the actual inner diameter, outer diameter, and wall thickness were smaller than designed (Figure 2Bi and 2Bii). The average outer diameter for each device ranged from 2.48 to 2.73 mm, a difference of 250 μm and the average inner diameter ranged from 0.76 to 0.81 mm, a difference of 40 μm. The difference can be attributed to the measurement technique (described in the Methods section). Additionally, standard deviations can be attributed to the measurement technique and inherent differences that occur in 3D printed devices. However, ultimately it is not critical to minimize the standard deviation across hydrogel rings because when an assay is performed with the rings the measurement made is the percent change in outer and inner diameter or area over time.

The addition of endothelial cells to line the lumen allows our method to further model the structure of a human blood vessel. Method 1 of creating collagen rings was adapted to make the post removable, enabling an endothelial cell suspension to be added to the lumen after gelation of the hydrogel ring (Figure 3). The post was removed and an endothelial cell suspension was pipetted into the lumen, after which the device was incubated for 1 hour. During this incubation period, the device was rotated 180 degrees every 5 minutes for the first 20 minutes and then every 10 minutes for an additional 40 minutes to ensure distribution of cells on all sides of the lumen. The spacer was removed and the Teflon tube was pushed down to reveal the ring with endothelial cells seeded in the lumen. The cells were then incubated in a well plate with cell culture media for 24 hours after which a live/dead stain was performed. Endothelial cells were viable at 24 hours and were evenly distributed within the lumen (Figure 3B). Imaging the hydrogel ring due to its free-standing nature can be challenging as it is suspended in liquid media. To increase the usability of this method, future work includes adding a holder to stabilize the ring without restricting its movement in three dimensions.

**Figure 3.**
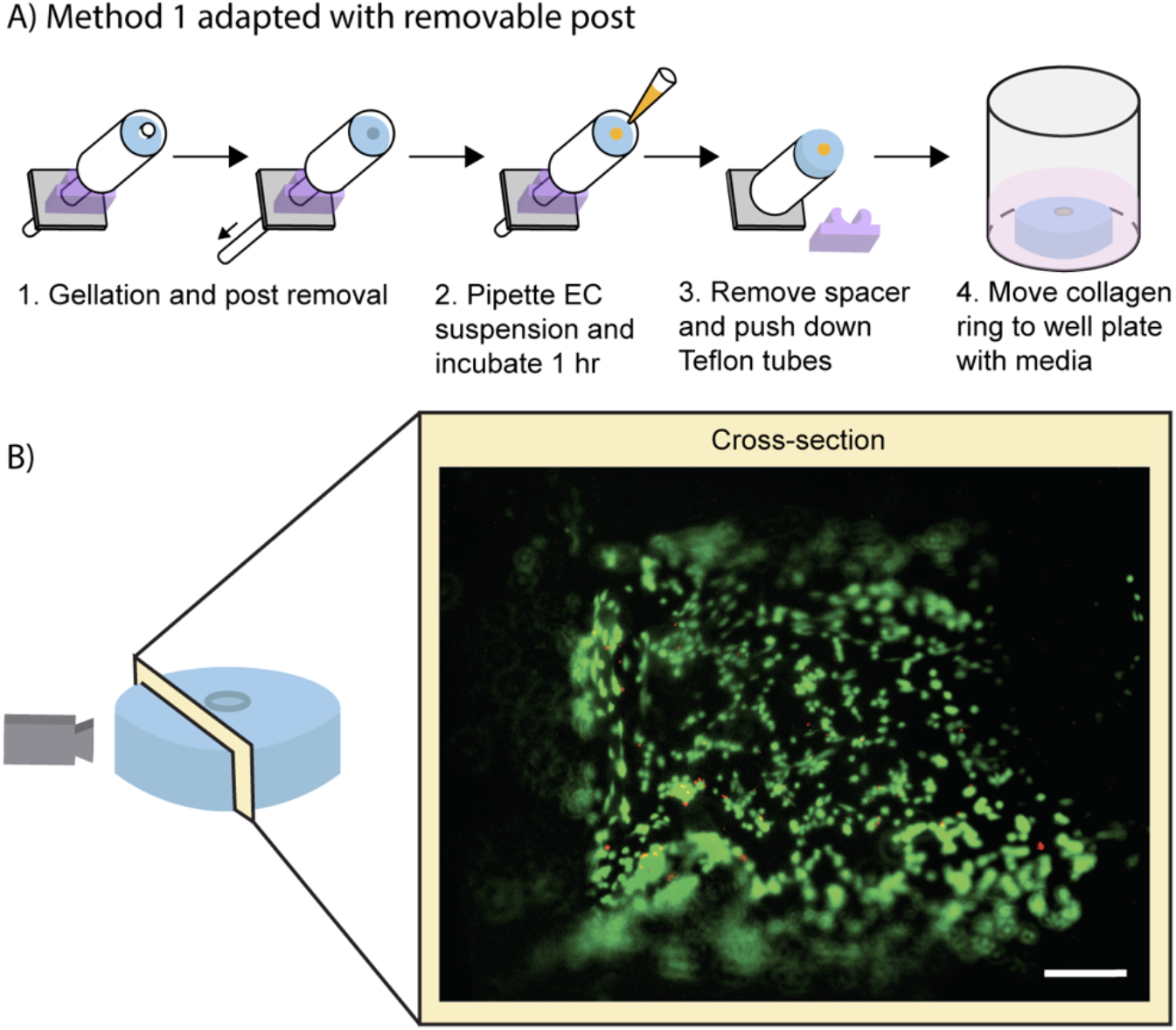
Addition of endothelial cells to the lumen of collagen I rings. (A) Workflow for adding the endothelial cell suspension to the gelled collagen rings after adapting Method 1. (B) Image of the lumen was taken after 24 hours of cell seeding to show the viability of the endothelial cells. Cells are stained with calcein AM (live, green) and ethidium-homodimer I (dead, red). Scale bar is 100 μm.

While Method 1 (Figure 1) works well for arrayed generation of hydrogel rings, we also sought to develop a method to make individual hydrogel rings with a simpler setup, avoiding the need to use F127 and multiple pieces (spacer, Teflon tubes, etc.). Method 2 uses a commercially available syringe that has been adapted to form collagen rings by cutting off the tip of the syringe and inserting a 3D printed core. Collagen laden with smooth muscle cells was then pipetted into the syringe and incubated to allow the collagen to gel. The 3D printed core was then pushed upwards (Figure 4A), revealing the collagen ring, and the ring was transferred to a well plate with cell culture media for storage. Following 5 days in culture, cell-laden hydrogel rings were submerged in a buffer (Tyrode’s Solution) and treated with either a control (additional Tyrode’s Solution) or a vasodilator (fasudil). The collagen rings were recorded for 20 minutes using a stereoscope to monitor any change in geometry and size. This imaging setup is cost effective (<$500 USD) and does not require specialized training to record the collagen rings. Images obtained from these recordings were analyzed using ImageJ, a free imaging processing software, to determine the change in percent area of the rings over time (Figure 4, FigureS1). A significant difference in ring area was observed when hydrogel rings were treated with a control (buffer) compared to the vasodilator (fasudil). We note that as the cells interact with the hydrogel, the mechanical stress experienced by the cells offers cues for cellular behavior and thus may affect the overall size change. ^24, 25^

**Figure 4.**
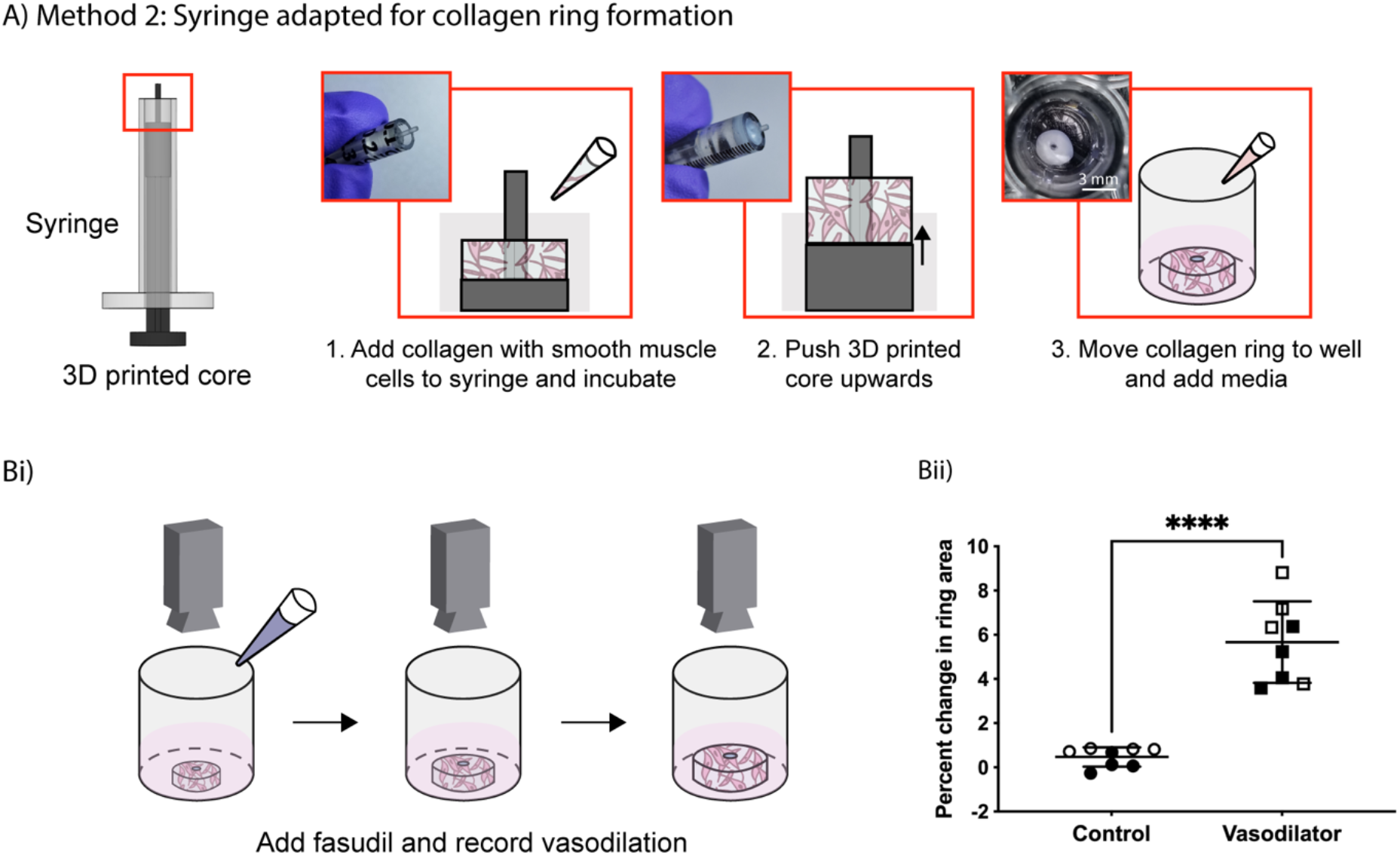
Percent change in ring area when human umbilical artery smooth muscle cells seeded in hydrogel rings were treated with a vasodilator (fasudil). (A) Method 2 workflow using a commercially available syringe with a 3D printed insertable core that has been adapted to make cell laden collagen rings. (Bi) The hydrogel rings were recorded for 20 minutes after addition of fasudil, and their percent change in area was calculated using ImageJ. (Bii) Percent change in ring area data for hydrogel rings treated with buffer (control) or vasodilator (fasudil). Data points are from 8 rings across 2 independent experiments; error bars are mean ± SD. A two-sample unpaired t-test (two-tailed) was used. ****p<0.0001.

## Methods

### Fabrication of device to mold hydrogel rings

#### Method 1 (Figures 1–3)

Molds were designed using Solidworks (Dassault Systems, Waltham, MA). Hydrogel ring molds were fabricated using a Form 2 SLA 3D printer (Formlabs, Somerville, MA) using Clear V4 resin (Formlabs) with a Z resolution of 0.05 mm. After printing, molds were cleaned in a FormWash with isopropyl alcohol (IPA) for 10 min, followed by a second wash with fresh IPA for 10 min. Molds were then dried using compressed air and cured under UV (FormCure, Formlabs) for 30 min at 60°C. A 3.5 mm thick polydimethylsiloxane (PDMS) spacer was made by milling a polystyrene mold (Datron Neo) and pouring PDMS (Sylgard^™^ 184, Dow) in a 1:8 ratio. The PDMS was left to cure at room temperature for 48 h. Prior to use for cell culture, molds were sprayed with 70% ethanol, air dried in a biosafety cabinet, and irradiated with UV light for 15 minutes.

#### Method 2 (Figure 4)

Cores were prepared using the same printing and sterilization protocols as Method 1. After posts were UV sterilized in the biosafety cabinet for 20 min, posts were soaked in 1% bovine serum albumin (BSA) for 40 minutes to prevent the collagen from sticking to the post, enabling easier removal of the hydrogel ring, and left to air dry before use. 1 mL syringes (COVIDIEN^™^) were trimmed at the 0.8 mL mark and the core was replaced with the 3D-printed posts.

### Cell culture of human umbilical artery smooth muscle cells (HUASMCs) and human umbilical vein endothelial cells (HUVECs)

Human umbilical artery smooth muscle cells (HUASMCs; Cell Applications, Inc) were cultured in human smooth muscle cell growth medium (SmGM; Cell Applications, Inc) supplemented with penicillin (100 units mL^-1^) and streptomycin (100 μg mL^-1^). Human umbilical vein endothelial cells (HUVECs; Lonza) were cultured in endothelial cell growth medium (EGM-2; Lonza) supplemented with penicillin (100 units mL^-1^) and streptomycin (100 μg mL^-1^). Culture flasks were maintained at 37 °C with 5% CO_2_. HUASMCs between passages 4 and 8 were used; HUVECs between passages 5 and 11 were used.

### Preparation of collagen I solution

HEPES buffer was prepared as 500 mM in 10X PBS and adjusted to a pH of 7.4 using NaOH pellets. The HEPES buffer was then thoroughly mixed with high concentration collagen I (8 – 10 mg mL^-1^; Corning) in a 1:9 ratio of HEPES to collagen.

### Encapsulation of HUASMCs in collagen I hydrogel

HUASMCs were trypsinized and resuspended at a concentration of 5 x 10^6^ cells mL^-1^ in growth medium (SmGM). The cell suspension was then added to the neutralized collagen solution for a final concentration of 1 x 10^6^ cells mL^-1^ and 6 mg mL^-1^ collagen I.

### Hydrogel ring fabrication

#### Method 1 (Figures 1–3)

Teflon tubing (McMaster Carr) with an inner diameter of 3.175 mm and an outer diameter of 6.35 mm was cut to 1 cm in length, sonicated in with 70% ethanol for 30 min, air dried, and then transferred onto the 3D printed molds with PDMS spacers. A 20% solution of Pluronic® F127 (P2443–1KG; referred to as F127) (Sigma Aldrich) was pipetted into the mold for the rings and then removed, leaving a thin layer of F127 and filling any gaps between the 3D printed device and the Teflon tubing. The devices were placed in a box with Kimwipes saturated in 1X PBS to prevent evaporation of any hydrogel. The box was placed in a 37 °C incubator for 5 min to fully set the F127. A tube of collagen was warmed up to room temperature to prevent it from dissolving the F127. The collagen was then pipetted into the molds, the molds placed back in the box with PBS, and incubated for 1 h. To remove the collagen rings the PDMS spacer was removed and the device was submerged in PBS. The Teflon tube was pushed down and the collagen rings were gently removed from the 3D printed post using tweezers or by gently shaking the device. They were then transferred to a 24 well plate with 500 μL of PBS in each well.

#### Method 2 (Figure 4)

For Method 2 the trimmed syringes and BSA soaked 3D printed cores were combined to form the mold. The cell-laden collagen solution was then carefully pipetted into the mold to avoid bubbles; depending on the tubing size, roughly 20 – 30 μL of cell-laden collagen solution was used per ring. After addition of hydrogel mixture, the molds were placed in a BioAssay Dish that was lined with Kimwipes soaked in 1X PBS. The BioAssay Dish containing molds was then transferred to an incubator to allow the collagen to gel at 37 °C. For the first 10 min, the BioAssay Dish was carefully flipped 180° every 2 min to ensure distribution of cells throughout the collagen.

After 1.5 h of incubation, the BioAssay Dish containing hydrogel ring molds was removed from the incubator and brought into a biosafety cabinet. The Teflon tubing was lowered and the hydrogel rings were transferred to individual wells in a 96-well plate. 100 μL of cell culture media was added to each well, and the surrounding wells were filled with 100 μL of 1X PBS to prevent evaporation of media. The hydrogel rings were then incubated at 37 °C with 5% CO2 until experimentation, with growth media replaced every 24 h.

### Measurement of hydrogel rings

Images were obtained using an Amscope MU1403B High Speed Microscope Camera mounted on an Amscope SM-3TZ-80S stereoscope (Amscope, Irvine, CA) and processed in FIJI (ImageJ). All images were thresholded between 40 and 255 prior to obtaining measurements. The outer diameter was measured manually by drawing a line across the image at 0, 45, 90, and 135 degrees and averaging those values; the same procedure was used for the inner diameter.

### Addition of endothelial cells to hydrogel rings (Figure 3)

Using Method I, a 3D printed mold was generated with a hole rather than a post. Teflon tubing (McMaster Carr) was run through each mold to act as a retractable post. Collagen I was added to the mold carefully to avoid bubbles and incubated for 1 h, as outlined above. After 1 h of incubation, the BioAssay Dish containing hydrogel ring molds was removed from the incubator and brought into a biosafety cabinet. The retractable Teflon tubing was then drawn down to reveal a lumen while the hydrogel rings remained in the mold. Approximately 5 μL solution of HUVECs (1 x 10^6^ cells mL^-1^) was carefully pipetted into the lumen for each ring. The devices were rotated 180 degrees every 5 min for the first 20 min and then 180 degrees every 10 min for the following 40 min to ensure even distribution of HUVECs within the lumen. After 1 h of incubation and rotation, hydrogel rings were removed in a solution of PBS and transferred to a well plate as described above. Each ring was incubated in 60 μL of EGM-2 media for 24 h. At the 24 hour mark rings were washed with PBS and stained with calcein AM (C3100MP, Fisher Scientific) and ethidium-homodimer 1 (E1169, Fisher Scientific) for 30 min at 37 °C, rinsed with PBS, and imaged using a fluorescence microscope (Zeiss Axiovert 200 and an Axiocam 503 mono camera; Carl Zeiss AG, Oberkochen, Germany).

### Dilation experiments

After 5 days in culture, cell-laden hydrogel rings were used for vasodilation experiments. Two hydrogel rings were transferred to individual wells of a new 96-well plate, and submerged in 60 μL Tyrode’s Solution (T2397, Sigma-Aldrich). After 5 min of equilibration the solution was removed and replaced with fresh Tyrode’s Solution. 6 μL of either additional Tyrode’s Solution (control) or fasudil (vasodilator) (HA-1077 dihydrochloride, Sigma-Aldrich) was then added. The well plate was then placed under a stereoscope and recorded for 20 min after which hydrogel rings were transferred to a new well plate containing cell media for staining and imaging. This process was repeated with additional hydrogel rings until the desired number of replicates had been tested.

### Imaging of vasodilation experiments and processing

Top-view images of hydrogel rings were recorded using an Amscope MU1403B High Speed Microscope Camera mounted on an Amscope SM-3TZ-80S stereoscope. Stills were obtained from video recordings for every 60 s (i.e., 21 stills were obtained for a 20 min recording). These images were then processed using FIJI (ImageJ) to calculate the total hydrogel area.

## Conclusion

We demonstrate that we have created a simple method for forming freestanding cell-laden hydrogel rings that can be used in a quantifiable assay for measuring dynamic change after treatment with a known vasodilator. Our goal was to create a method that does not rely on animal models or rigid devices to stabilize the lumen. Our method takes the unique approach to being freestanding which allows unrestricted movement in all three dimensions in response to vasoactive compounds. This method enables researchers to study the response of a controlled human cell-laden hydrogel ring to different inflammatory drugs or molecules, as well as study the cellular communication that is important to understanding the pathways that are activated during inflammation. So far, we have only tested the model with fasudil; additional vasodilators and vasoconstrictors would need to be tested to further validate our method for use in vasodilator and vasoconstrictor studies. Additionally, alignment of the smooth muscle cells was not achieved in this current work. This will be a focus of future work to ensure smooth muscle cells are able to constrict and dilate while in the alignment observed in blood vessels. In future work we will also work to achieve lumen size and wall thickness relevant to smaller vessels in the human body. We anticipate this can be addressed using higher resolution fabrication methods. Additionally, we will integrate these freestanding lumens with other co- and multiculture platforms to incorporate fibroblasts, smooth muscle cells, endothelial cells, and other stromal cells important for modeling disease-specific vasodilation, as well as cell types in circulating blood including leukocytes that may interact with endothelial cells and be recruited to neighboring tissue. We envision that our model will be important for studying a range of diseases such as asthma, cancer, autoimmune disease, and other diseases that have periods of induced inflammation, vasodilation, or constriction.

## Acknowledgements

This work was supported by the University of Washington, National Institutes of Health (NIH1R35GM128648, ABT, EB, AMD, HGL, TLVN), the Society for Laboratory Automation and Screening (SLASFG2020, UNL), the Howard Hughes Medical Institute James H. Gilliam Fellowship for Advanced Study program (GT14938, TLVN), Washington Research Foundation Fellowship (Washington Research Foundation, HGL), CoMotion Mary Gates Innovation Scholars Fellowship (University of Washington CoMotion, HGL). Any opinions, findings, and conclusions or recommendations expressed in this material are those of the author(s) and do not necessarily reflect those of the Society for Laboratory Automation and Screening or the NIH.

