## Supplementary Information for "Freestanding hydrogel lumens for modeling blood vessels and vasodilation"

### Freestanding hydrogel lumens for biological assays

[1]Department of Chemistry, University of Washington, Seattle, WA, USA
[2]Department of Urology, University of Washington School of Medicine, Seattle, WA, USA
*Denotes co-first author

Contents:
Table S1: Comparison of different methods
Table S2: Full set of data for Figure 2Bi and 2Bii
Figure S1: Images of collagen rings during vasodilation experiment

Table S1. Comparison of general features of excised animal tissue models, standard closed microfluidic lumen models, and our freestanding method

| | Excised Animal Tissue[1,2] | Standard Closed Channel Microfluidic Lumen Models[3,4] | Our Freestanding Method |
|---|---|---|---|
| Freestanding | Yes | No | Yes |
| Requires Animal Tissue | Yes | No | No |
| Tissue Layers Are Possible | Yes | Yes | Yes |
| Human Cells Can Be Incorporated | No | Yes | Yes |

Table S2. Full set of data for Figure 2Bi and 2Bii. Each ring was measured four times for outer diameter (OD) and inner diameter (ID).

| | Length (mm) | Average (mm) | SD (mm) |
|---|---|---|---|
| Expt 1-rack 1_ring 1-OD | 2.568 | 2.591 | 0.074 |
| | 2.558 | | |
| | 2.700 | | |
| | 2.536 | | |
| Expt 1-rack 1_ring 1-ID | 0.854 | 0.828 | 0.018 |
| | 0.815 | | |
| | 0.828 | | |
| | 0.813 | | |
| Expt 1-rack 1_ring 2-OD | 2.508 | 2.590 | 0.062 |
| | 2.654 | | |
| | 2.615 | | |
| | 2.584 | | |
| Expt 1-rack 1_ring 2-ID | 0.807 | 0.790 | 0.027 |
| | 0.807 | | |
| | 0.794 | | |
| | 0.750 | | |
| Expt 1-rack 1_ring 3-OD | 2.280 | 2.452 | 0.140 |
| | 2.502 | | |
| | 2.612 | | |
| | 2.414 | | |

| | | | |
|---|---|---|---|
| Expt 1-rack 1_ring 3-ID | 0.835 | 0.823 | 0.016 |
| | 0.823 | | |
| | 0.833 | | |
| | 0.800 | | |
| Expt 1-rack 1_ring 4-OD | 2.368 | 2.445 | 0.068 |
| | 2.463 | | |
| | 2.420 | | |
| | 2.528 | | |
| Expt 1-rack 1_ring 4-ID | 0.749 | 0.765 | 0.021 |
| | 0.796 | | |
| | 0.753 | | |
| | 0.761 | | |
| Expt 1-rack 1_ring 5-OD | 2.621 | 2.610 | 0.089 |
| | 2.642 | | |
| | 2.692 | | |
| | 2.483 | | |
| Expt 1-rack 1_ring 5-ID | 0.733 | 0.728 | 0.062 |
| | 0.686 | | |
| | 0.813 | | |
| | 0.679 | | |
| Expt 1-rack 1_ring 6-OD | 2.258 | 2.164 | 0.152 |
| | 1.938 | | |
| | 2.204 | | |
| | 2.254 | | |
| Expt 1-rack 1_ring 6-ID | 0.807 | 0.810 | 0.029 |
| | 0.799 | | |
| | 0.783 | | |
| | 0.850 | | |
| Expt 1-rack 2_ring 1-OD | 2.725 | 2.713 | 0.033 |
| | 2.678 | | |
| | 2.696 | | |
| | 2.752 | | |
| Expt 1-rack 2_ring 1-ID | 0.771 | 0.763 | 0.033 |
| | 0.736 | | |
| | 0.738 | | |
| | 0.805 | | |

| | | | |
|---|---|---|---|
| Expt 1-rack 2_ring 2-OD | 2.674 | 2.659 | 0.069 |
| | 2.674 | | |
| | 2.724 | | |
| | 2.562 | | |
| Expt 1-rack 2_ring 2-ID | 0.874 | 0.761 | 0.096 |
| | 0.668 | | |
| | 0.696 | | |
| | 0.805 | | |
| Expt 1-rack 2_ring 3-OD | 2.456 | 2.429 | 0.069 |
| | 2.504 | | |
| | 2.414 | | |
| | 2.341 | | |
| Expt 1-rack 2_ring 3-ID | 0.846 | 0.810 | 0.043 |
| | 0.787 | | |
| | 0.845 | | |
| | 0.761 | | |
| Expt 1-rack 2_ring 4-OD | 2.589 | 2.554 | 0.068 |
| | 2.452 | | |
| | 2.580 | | |
| | 2.595 | | |
| Expt 1-rack 2_ring 4-ID | 0.870 | 0.840 | 0.020 |
| | 0.833 | | |
| | 0.828 | | |
| | 0.828 | | |
| Expt 1-rack 2_ring 5-OD | 2.684 | 2.691 | 0.023 |
| | 2.716 | | |
| | 2.662 | | |
| | 2.700 | | |
| Expt 1-rack 2_ring 5-ID | 0.897 | 0.888 | 0.028 |
| | 0.918 | | |
| | 0.887 | | |
| | 0.850 | | |
| Expt 1-rack 2_ring 6-OD | 2.684 | 2.726 | 0.036 |
| | 2.769 | | |
| | 2.715 | | |
| | 2.737 | | |

| | | | |
|---|---|---|---|
| Expt 1-rack 2_ring 6-ID | 0.807 | 0.783 | 0.023 |
| | 0.765 | | |
| | 0.798 | | |
| | 0.761 | | |
| Expt 2-rack 1_ring 1-OD | 2.753 | 2.733 | 0.031 |
| | 2.697 | | |
| | 2.718 | | |
| | 2.763 | | |
| Expt 2-rack 1_ring 1-ID | 0.785 | 0.786 | 0.021 |
| | 0.815 | | |
| | 0.777 | | |
| | 0.766 | | |
| Expt 2-rack 1_ring 2-OD | 2.742 | 2.701 | 0.034 |
| | 2.668 | | |
| | 2.677 | | |
| | 2.715 | | |
| Expt 2-rack 1_ring 2-ID | 0.759 | 0.724 | 0.043 |
| | 0.675 | | |
| | 0.701 | | |
| | 0.761 | | |
| Expt 2-rack 1_ring 3-OD | 2.457 | 2.402 | 0.063 |
| | 2.363 | | |
| | 2.334 | | |
| | 2.454 | | |
| Expt 2-rack 1_ring 3-ID | 0.812 | 0.720 | 0.066 |
| | 0.659 | | |
| | 0.716 | | |
| | 0.694 | | |
| Expt 2-rack 1_ring 4-OD | 2.758 | 2.720 | 0.037 |
| | 2.742 | | |
| | 2.677 | | |
| | 2.704 | | |
| Expt 2-rack 1_ring 4-ID | 0.828 | 0.806 | 0.033 |
| | 0.833 | | |
| | 0.761 | | |
| | 0.802 | | |

| | | | |
|---|---|---|---|
| Expt 2-rack 1_ring 5-OD | 2.773 | 2.830 | 0.062 |
| | 2.868 | | |
| | 2.897 | | |
| | 2.780 | | |
| Expt 2-rack 1_ring 5-ID | 0.692 | 0.719 | 0.056 |
| | 0.759 | | |
| | 0.772 | | |
| | 0.654 | | |
| Expt 2-rack 1_ring 6-OD | 2.630 | 2.578 | 0.075 |
| | 2.466 | | |
| | 2.607 | | |
| | 2.609 | | |
| Expt 2-rack 1_ring 6-ID | 0.775 | 0.822 | 0.041 |
| | 0.842 | | |
| | 0.805 | | |
| | 0.867 | | |
| Expt 2-rack 2_ring 1-OD | 2.758 | 2.719 | 0.069 |
| | 2.621 | | |
| | 2.722 | | |
| | 2.774 | | |
| Expt 2-rack 2_ring 1-ID | 0.823 | 0.843 | 0.018 |
| | 0.865 | | |
| | 0.835 | | |
| | 0.850 | | |
| Expt 2-rack 2_ring 2-OD | 2.732 | 2.755 | 0.016 |
| | 2.763 | | |
| | 2.767 | | |
| | 2.756 | | |
| Expt 2-rack 2_ring 2-ID | 0.825 | 0.824 | 0.008 |
| | 0.817 | | |
| | 0.820 | | |
| | 0.835 | | |
| Expt 2-rack 2_ring 3-OD | 2.774 | 2.825 | 0.088 |
| | 2.951 | | |
| | 2.756 | | |
| | 2.819 | | |

| | | | |
|---|---|---|---|
| Expt 2-rack 2_ring 3-ID | 0.754 | 0.754 | 0.043 |
| | 0.701 | | |
| | 0.805 | | |
| | 0.757 | | |
| Expt 2-rack 2_ring 4-OD | 2.761 | 2.758 | 0.003 |
| | 2.757 | | |
| | 2.755 | | |
| | 2.758 | | |
| Expt 2-rack 2_ring 4-ID | 0.787 | 0.769 | 0.022 |
| | 0.767 | | |
| | 0.738 | | |
| | 0.783 | | |
| Expt 2-rack 2_ring 5-OD | 2.725 | 2.726 | 0.025 |
| | 2.697 | | |
| | 2.724 | | |
| | 2.758 | | |
| Expt 2-rack 2_ring 5-ID | 0.831 | 0.803 | 0.038 |
| | 0.759 | | |
| | 0.783 | | |
| | 0.839 | | |
| Expt 2-rack 2_ring 6-OD | 2.457 | 2.584 | 0.098 |
| | 2.687 | | |
| | 2.566 | | |
| | 2.625 | | |
| Expt 2-rack 2_ring 6-ID | 0.664 | 0.720 | 0.044 |
| | 0.738 | | |
| | 0.768 | | |
| | 0.709 | | |

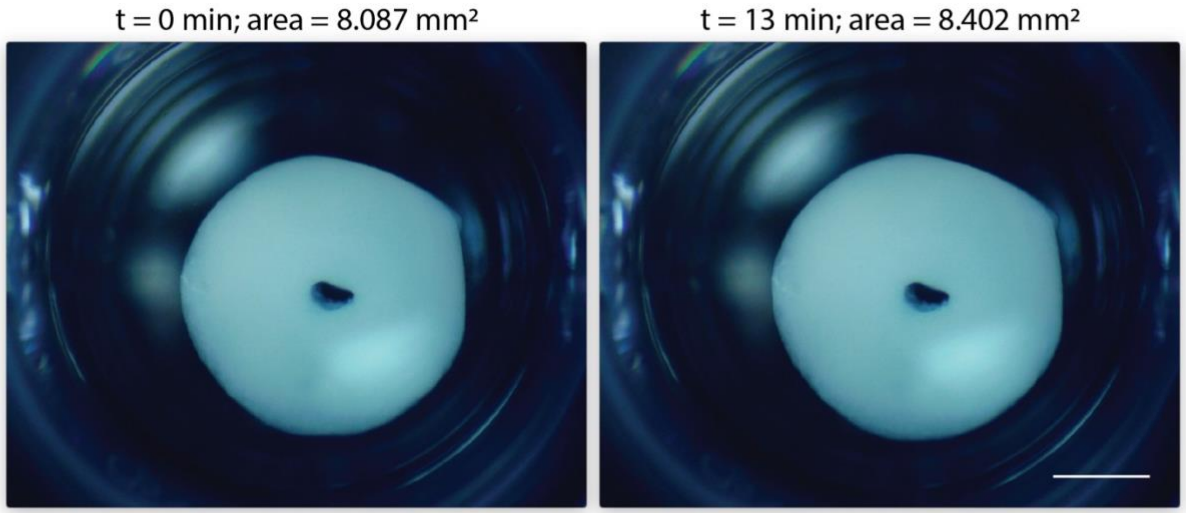


Figure S1: Images of smooth muscle cell laden collagen I rings during experimentation with a vasodilator (fasudil). Images are at t = 0, addition of fasudil, and at t = 13 min. Change in ring area is not visible by eye; it is quantifiable using image processing software (see methods section)
